# Contribution of front-line, standard of care drugs to bactericidal responses, resistance emergence, and cure in murine models of easy- or hard-to-treat tuberculosis disease

**DOI:** 10.1101/2024.12.18.629278

**Authors:** Nathan Peroutka-Bigus, Elizabeth J. Brooks, Michelle E. Ramey, Hope D’Erasmo, Jacqueline P. Ernest, Allison A. Bauman, Lisa K Woolhiser, Radojka M. Savic, Anne J. Lenaerts, Bree B. Aldridge, Jansy P. Sarathy, Gregory T. Robertson

**Affiliations:** Mycobacteria Research Laboratories, Department of Microbiology, Immunology and Pathology, Colorado State University, Fort Collins, Colorado, USA; Department of Molecular Biology and Microbiology, Tufts University School of Medicine, Boston, MA, USA; Department of Bioengineering and Therapeutic Sciences, University of California San Francisco, San Francisco, CA, USA; The Stuart B. Levy Center for Integrated Management of Antimicrobial Resistance, Boston, MA, USA; Department of Biomedical Engineering, Tufts University School of Engineering, Medford, MA, USA; Center for Discovery and Innovation, Hackensack Meridian Health, Nutley, NJ, USA; Hackensack Meridian School of Medicine, Department of Medical Sciences, Nutley, NJ, USA

**Keywords:** Tuberculosis, relapse, caseum, rifafour, C3HeB/FeJ

## Abstract

By assessing the standard of care regimen for tuberculosis (TB) in BALB/c and C3HeB/FeJ mice, we demonstrate that rifampin, with or without pyrazinamide, is essential to effective bactericidal response and resistance suppression. Potency measurements in an *in vitro* lipid-rich model and rabbit caseum assay recapitulate the significance of rifampin as a sterilizing agent. These outcomes align with clinical performance, thus emphasizing the value of *in vitro* predictive tools and murine TB models with human-like pathology.

---

Wallace Fox described human clinical studies that led to the current standard-of-care chemotherapy for tuberculosis (1). Rifampin (R) and pyrazinamide (Z) were denoted as key sterilizing drugs, with isoniazid (H) contributing to bactericidal responses, while ethambutol (E), a bacteriostatic drug, was described as contributing little to bactericidal responses or sterilizing cure (1).

Using two pathologically distinct murine tuberculosis (TB) models we sought to interrogate treatment outcomes for the standard-of-care regimen (see **Methods, supplemental data**). In BALB/c mice chronically infected with *Mycobacterium tuberculosi*s (Mtb) Erdman [without the caseating granulomas seen in patients (2)], HRZE reduced BALB/c lung burdens by 3.28 logs after 1-month treatment, by 5.60 logs after 2-months (3/5 mice remained culture positive) and returned no CFU after 3-months of 2HRZE/HR (**Fig. 1A-D, Table S1**), indicating potent bactericidal responses. HRZE was equally potent in C3HeB/FeJ mice chronically infected with Mtb Erdman [showing more human-like, heterogeneous lung pathology (2-4)], reducing lung burdens by 4.97 logs after 1-month treatment, by 6.54 logs after 2-months (4/6 mice remained culture positive) (**Fig. 1E-H, Table S1**), and returned no CFU after 3-months of 2HRZE/HR in C3HeB/FeJ except for 1 mouse returning a single CFU. These results agree with previously published findings (5).

**FIG 1.**
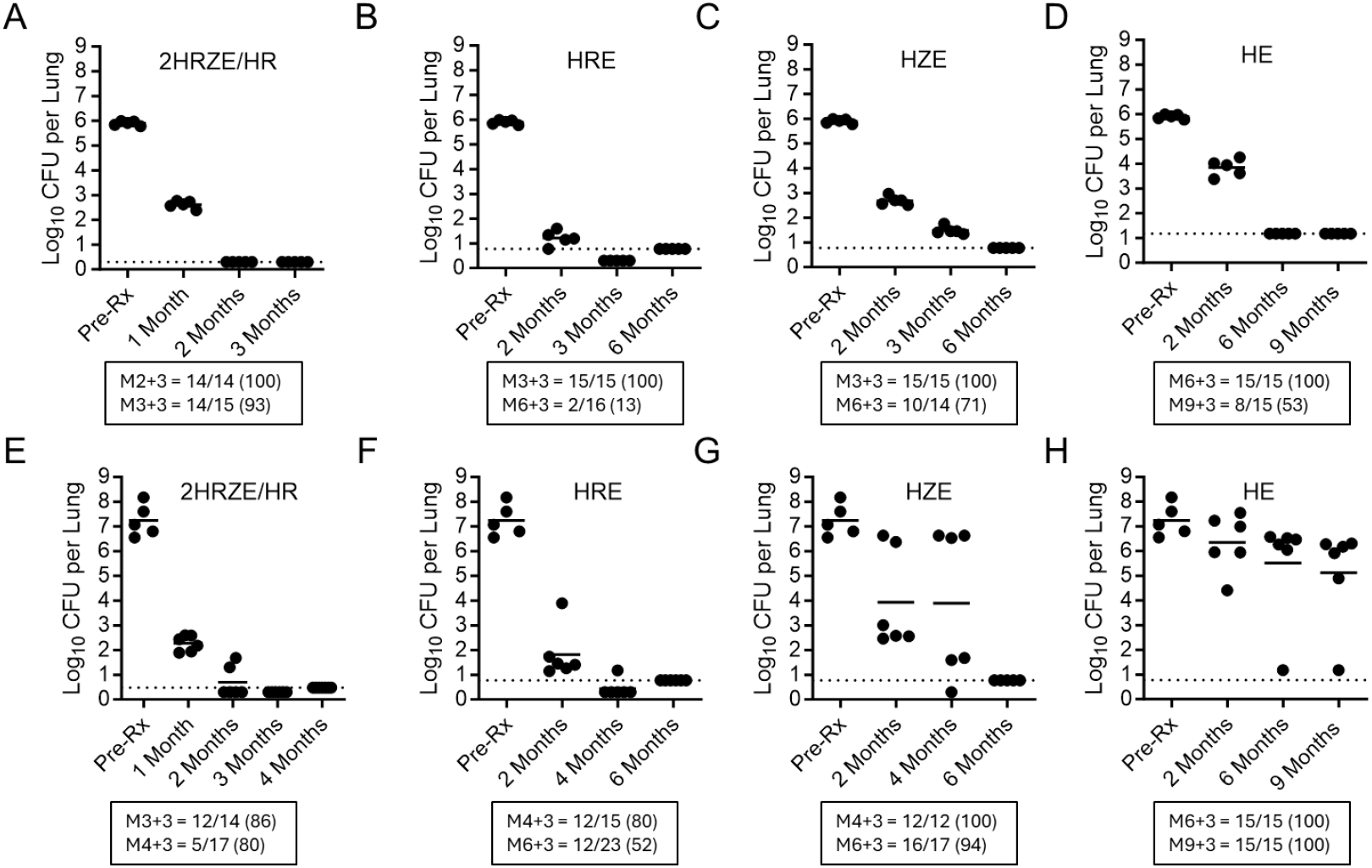
*Mycobacterium tuberculosis* Erdman CFU burdens in lungs of BALB/c (A-D) and C3HeB/FeJ (E-H) mice at the start of treatment (Pre-Rx) or after treatment for the indicated number of months with 2HRZE/HR (A, E), HRE (B, F), HZE (C, G), or HE (D, H). Closed circles represent individual mice. The solid horizontal line is the group mean value. The dashed horizontal line is the upper lower limit of detection. The box beneath represents the number of mice that relapsed 3 months after the indicated treatment duration over the group total; the proportion of mice relapsing is listed in parentheses.

HRE, HZE, and HE proved significantly less bactericidal versus HRZE in BALB/c mice, promoting 4.69, 3.21, 2.06 log10 CFU reductions in lungs, respectively, after 2-months treatment (highest P value = 0.0001). No CFU were cultured after 6 months HRE or HZE, and only one of 5 mice in the HE group was culture positive. All HE-treated mice were culture negative by month 9 (**Figs. 1B-1D, Table S1**).

Similar to the BALB/c arm, HRE reduced C3HeB/FeJ lung burdens by 5.43 logs after 2-months treatment (**Fig. 1F, Table S1**), revealing a minor role for Z in the HRZE bactericidal response. However, substitution of Z for R in HZE (HZE vs HRE, P = 0.0482) or adding R or Z to HE (HE vs HRE, P < 0.0001; HE vs HZE, P = 0.0214), improved efficacy significantly [3.31 and 0.89 logs from the start of treatment, respectively] (**Figs. 1E-1H, Table S1**). Interestingly, a bi-modal response was seen in C3HeB/FeJ administered regimens lacking R or RZ; whereby a subpopulation of mice was less responsive to drug treatment based on pulmonary pathology (10-12). This was not similarly observed in the BALB/c arm. Only in combinations including R or RZ, was a more effective and uniform treatment response observed in C3HeB/FeJ mice, similar to the BALB/c arm (compare **Figs. 1C-1D** to **Figs. 1G-1H**). As described by Fox (1), more mice underwent relapse following 6 months of treatment with regimens lacking R or RZ. HE was the least effective in C3HeB/FeJ, with all mice relapsing following 9-months treatment (**Fig. 1H, Tables S2A, B**).

Co-plating on antibiotic-containing agar plates after 2HRZE/HR therapy in C3HeB/FeJ resulted in rare resistant isolates to H, R, or E (see **Table S3A**), similar to the HRE or HZE groups. In contrast, 5/6 C3HeB/FeJ mice treated with HE had a high number of isolates that grew on 0.2 mg/L of H, with a low frequency of resistance to E suggesting that H resistance occurred without loss of susceptibility to E in many cases (see **Table S3A**). Higher rates of resistance were observed in the C3HeB/FeJ study arm during relapse, which was far lower in the BALB/c arm (see **Tables S3B** and **S4B**).

To further investigate why R was a good partner drug *in vivo*, we systematically evaluated *in vitro* drug combination effects of six drug pairs from HRZE using data from Larkins-Ford et al (6). Pairwise potencies were evaluated in a lipid-rich environment (butyrate), where Z shows activity, using the infinite growth rate (GRinf), a metric of the combination potency and predictor of relapsing outcomes (6). HR and RE were the most potent, and more potent than HZ and ZE (**Figure 2**). Together, this data suggested that R pairs well with E and H in lipid-rich environments.

**Figure 2.**
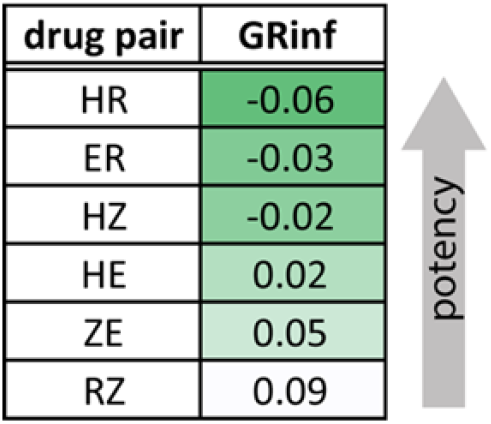
Pairwise drug combination potencies in medium with butyrate as the sole carbon source measured as the GRinf after 6d of treatment using diagonal measurement of n-way drug interactions (DiaMOND). Data are derived from Larkins-Ford et al. (6).

HRZE was examined in *ex vivo* rabbit caseum (**Figure 3**), evaluating drug potency against non-replicating bacteria (7). Previously, H, Z and E were shown to be minimally active (casMBC_90_s > 512 µM). Only R produced significant killing (casMBC_90_ of 10 µM) (8, 9). Here, R concentrations were tested centered around 4 µM (0.0156 to 64 µM) to approximate the average concentrations in caseum (C_ave[0–24]_). H, Z and E were held static at 2, 56 and 8 µM, their respective caseum C_ave[0–24]_s (**Figure 3B**). Increased R exposure in HRZE produced increased bacterial killing in rabbit caseum, achieving 1-log killing when all drugs were present at C_ave[0–24]_ (datapoint 5) (**Figure 3A**). This result suggests a role for the potency of R against nonreplicating Mtb from caseous granulomas in driving treatment efficacy in C3HeB/FeJ mice.

**Figure 3.**
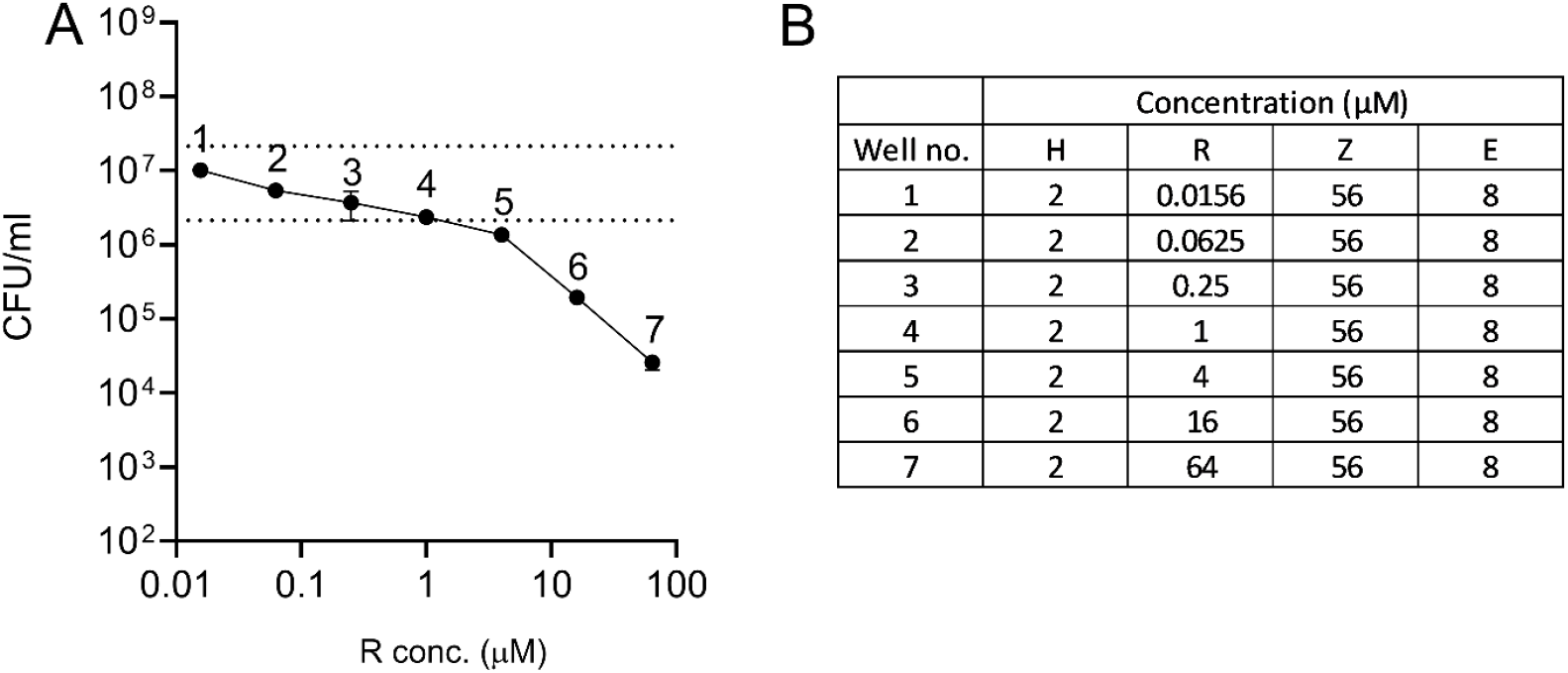
Bactericidal activity of HRZE in *ex vivo* rabbit caseum (A). Bacterial burden and R concentration are expressed on log scales. The dotted line indicates the bacterial burden in the DMSO-only control well and the cutoff for a 1-log reduction. Concentrations used for each drug at each datapoint are shown in the inset table (B). Data points and error bars represent the means and standard deviations for three technical replicates each.

In conclusion, while the regimens 2HRZE/HR and HRE showed nearly identical activities in BALB/c and C3HeB/FeJ mice, regimens lacking R or RZ were far less efficacious in C3HeB/FeJ mice in terms of bactericidal response, prevention of relapse, and suppression of resistance emergence. These results align well with observations by Fox (1), and with data reported in 8-week early bactericidal activity (EBA) trials (13), highlighting the contribution of R or RZ to the current front-line TB regimen (14). High rates of resistance were noted for C3HeB/FeJ given only HE, but not for mice on Z or RZ regimens. Fox similarly reported a minor role for E in initial resistance to H, and suppression of R-resistance in cases where the infection is resistant to H (1). Collectively this work illustrates the ability of diverse murine TB efficacy models of increasing complexity to better highlight differences in regimen behavior(s) and assessing contribution of individual drugs to regimens.

## Acknowledgments

This work was supported by the Bill and Melinda Gates Foundation under grant ID numbers INV-009105, ‘TB Drug Accelerator: TB mouse *in vivo* models’ [G.T.R.], OPP1033596, “Evaluation of a New Murine Model for Testing Tuberculosis Chemotherapy” and OPP1037174, “Qualification of C3HeB/FeJ mice for Experimental Chemotherapy of Tuberculosis” [A.J.L.]. INV-027276 “Design of combination therapies for tuberculosis” [B.B.A.]. We acknowledge Laura Via and Clifton Barry 3rd at the National Institute of Health for rabbit caseum, and the staff of the Laboratory Animal Resources at Colorado State University for animal care.

